# Estimates for quality of life loss due to RSV

**DOI:** 10.1101/321844

**Authors:** David Hodgson, Katherine E. Atkins, Marc Baguelin, Jasmina Panovska-Griffiths, Dominic Thorrington, Albert Jan van Hoek, Hongxin Zhao, Ellen Fragaszy, Andrew C. Hayward, Richard Pebody

**Affiliations:** Centre for Mathematics, Physics and Engineering in the Life Sciences and Experimental Biology, University College London, London, UK.; Clinical Operational Research Unit, Department of Mathematics, University College London, London, UK.; Centre for the Mathematical Modelling of Infectious Diseases, London School of Hygiene & Tropical Medicine, London, UK; Department of Infectious Disease Epidemiology, Faculty of Epidemiology and Population Health, London School of Hygiene & Tropical Medicine, London, UK; Respiratory Diseases Department, Public Health England, London, UK; department of Applied Health Research University College London, London, UK; Department of Epidemiology and Surveillance, National Institute for Public Health and Environment, Bilthoven, The Netherlands.; Centre for Public Health Data Science, Institute of Health Informatics, University College London, London, UK; Department of Epidemiology and Public Health, University College London, London, UK.

**Keywords:** Quality-adjusted life years, Respiratory disease, Cost-effectiveness, EQ-5D, Health-related quality of life, Respiratory Syncytial Virus, Human

## Abstract

A number of vaccines against Respiratory Syncytial Virus (RSV) infection are approaching licensure. Deciding which RSV vaccine strategy, if any, to introduce, will partly depend on cost-effectiveness analyses, which compares the relative costs and health benefits of a potential vaccination programme. Health benefits are usually measured in Quality Adjusted Life Year (QALY) loss, however, there are no QALY loss estimates for RSV that have been determined using standardised instruments. Moreover, in children under the age of five years in whom severe RSV episodes predominantly occur, there are no appropriate standardised instruments to estimate QALY loss. We estimated the QALY loss due to RSV across all ages by developing a novel regression model which predicts the QALY loss without the use of standardised instruments. To do this, we conducted a surveillance study which targeted confirmed episodes in children under the age of five years (confirmed cases) and their household members who experienced symptoms of RSV during the same time (suspected cases.) All participants were asked to complete questions regarding their health during the infection, with the suspected cases aged 5–14 and 15+ years old additionally providing Health-Related Quality of Life (HR-QoL) loss estimates through completing EQ-5D-3L-Y and EQ-5D-3L instruments respectively. The questionnaire responses from the suspected cases were used to calibrate the regression model. The calibrated regression model then used other questionnaire responses to predict the HR-QoL loss without the use of EQ-5D instruments. The age-specific QALY loss was then calculated by multiplying the HR-QoL loss on the worst day predicted from the regression model, with estimates for the duration of infection from the questionnaires and a scaling factoring for disease severity. Our regression model for predicting HR-QoL loss estimates that for the worst day of infection, suspected RSV cases in persons five years and older who do and do not seek healthcare have an HR-QoL loss of 0·616 (95% CI 0·155–1·371) and 0·405 (95% CI 0·111–1·137) respectively. This leads to a QALY loss per RSV episode of 1·950 × 10^−3^ (95% CI 0·185 × 10^−3^ –9·578 × 10^−3^) and 1·543 × 10^−3^ (95% CI 0·136 × 10^−3^ –6·406 × 10^−3^) respectively. For confirmed cases in a child under the age of five years who sought healthcare, our model predicted a HR-QoL loss on the worst day of infection of 0·820 (95% CI 0·222–1·450) resulting in a QALY loss per RSV episode of 3·823 × 10^−3^ (95% CI 0·492 × 10^−3^ –12·766 × 10^−3^). Combing these results with previous estimates of RSV burden in the UK, we estimate the annual QALY loss of healthcare seeking RSV episodes as 1,199 for individuals aged five years and over and 1,441 for individuals under five years old. The QALY loss due to an RSV episode is less than the QALY loss due to an Influenza episode. These results have important implications for potential RSV vaccination programmes, which has so far focused on preventing infections in infants—where the highest reported disease burden lies. Future potential RSV vaccination programmes should also evaluate their impact on older children and adults, where there is a substantial but unsurveilled QALY loss.

## Introduction

Respiratory Syncytial Virus (RSV) is a leading cause of lower respiratory tract infection in infants, accounting for three million hospital admissions and 60,000 deaths in children less than five years of age annually.^1,2^ Despite its health burden, there is no licensed vaccine for RSV, leaving infants vulnerable to infection. However, with over 50 RSV vaccine candidates currently in preclinical and clinical trials, it is likely that a vaccine will come to market in the near future.^3^ Decisions regarding the introduction of future vaccines will be informed by their projected impact and cost-effectiveness.

Cost-effectiveness analysis relies on the existence of measures for morbidity and mortality associated with RSV episodes expressed in terms of QALY loss.^4^ QALY loss is determined by considering the duration for which the RSV-specific loss of Health-Related Quality of Life (HR-QoL) is experienced. HR-QoL is evaluated through the use of standardised instruments, such as EuroQol’s EQ-5D that considers the physical, mental and emotional effects of an infection.^5^ The HR-QoL loss due to RSV across all ages through the use of standardised methods has yet to be performed. Moreover, these methods are not validated to capture the HR-QoL in very young children, in whom severe RSV episodes predominantly occur.^6^

In this study, we determined the QALY loss due to an RSV episode across all ages by conducting a surveillance study among recently confirmed RSV cases aged 0-4 years and suspected RSV cases in older household members. For both confirmed and suspected infections, we determined indicators of the severity of the RSV episode using questionnaires that evaluated loss in Visual Analogue Scale (VAS), lost school/work days (if appropriate), coughing severity and healthcare-seeking behaviour. For suspected cases in those aged 5-14 and 15+, we used EuroQoL EQ-5D-3L-Y^7–9^ and EQ-5D-3L^7,8^ questionnaires, respectively, to also determine the HR-QoL loss. Using these responses, we performed a regression analysis to evaluate the relationship between the HR-QoL loss and indicators of disease severity. In addition to estimating QALY loss stratified by healthcare-seeking behaviour and disease severity in older children and adults, our approach provides a novel method to calculate QALY loss for RSV episodes in young children.

## Methods

### Study recruitment

During the 2016-17 RSV season, confirmed cases of RSV in children under the age of 5 years from the previous two weeks were extracted on dates 13th December, 25th December 2016, and 3rd January 2017 from the Public Health England (PHE) Respiratory DataMart surveillance (RDMS) system.^10^ For all the confirmed cases for whom name, date of birth and National Health Service (NHS) number were provided, home addresses were obtained from the PHE Patient Demographic Service. For all obtained home addresses, a questionnaire pack addressed to the parent or guardian of the confirmed case, was sent the day after its extraction date from the PHE RDMS system. Each questionnaire pack consisted of three questionnaires, an information sheet, and a stamped addressed return envelope. The Index Questionnaire requested information about the recent RSV episode in the confirmed case. The other two questionnaires requested information about suspected RSV episodes in older household members; those aged 5-14 years (5-14 Questionnaire) and those aged 15 years or older (15+ Questionnaire). Suspected RSV cases were defined as persons who share a household with the confirmed case and who experienced an onset of RSV-like symptoms (runny or blocked nose, fever, coughing, and/or a sore throat) between five days before and five days after the onset of symptoms in the confirmed case.^11,12^ All questionnaire responses which answered at least one question were included in the data analysis. For the QALY estimate, we excluded responses which did not indicate a duration of coughing and those that indicated a duration of more than 22 days as these were extreme outliers in the sample (three times the upper bound of the interquartile range) in addition to being longer than the duration of infection for RSV observed in existing studies.^11^

### Questionnaire information

The Index Questionnaire was completed by a parent or guardian on behalf of the confirmed case, the 5-14 Questionnaire on behalf of or by the child themselves and the 15+ Questionnaire by the adolescent or adult. The Index Questionnaire requested information on (i) the age of the child, (ii) the confirmed case’s symptoms (runny/blocked nose, fever, coughing, sore throat), (iii) the healthcare seeking behaviour (no healthcare sought, healthcare sought), (iv) coughing severity (mild/no coughing, severe coughing) and (v) a Visual Analogue Scale (VAS) for the worst day of the recent infection and the day of questionnaire completion. A VAS was presented for health from 0 (worst health) to 100 (best health) for both days and the difference between the VAS scores was defined as the VAS score loss due to an RSV episode. In addition to the questions asked in the Index Questionnaire, the 5-14 and the 15+ Questionnaires also asked (vi) the time taken off school/work due to symptoms (productivity) and (vii) EuroQol EQ-5D-3L-Y^7–9^ or EQ-5D-3L questionnaires to determine Health-related Quality of Life weight at baseline and on the worst day of suspected RSV infection,^7,8^ respectively. See Appendix I for full questionnaire packs.

The EuroQol ED-5D-3L-Y^7–9^ and EQ-5D-3L^7,8^ questionnaires use a UK specific Time Trade-Off scoring tariff to determine the HR-QoL weight according to five dimensions: mobility, self-care, usual activities, pain/discomfort, and anxiety/depression. Respondents were asked to complete the EQ-5D responses for their health or the health of the child on the day they received the questionnaire (baseline) and for the worst day of infection. We refer to this HR-QoL weight on the worst day of infection as the raw maximum HR-QoL weight for an RSV episode and the difference in the HR-QoL weights between the baseline and the worst day of infection as the raw maximum HR-QoL loss.

### Regression model

EQ-5D instruments are not validated for children under five years of age so we cannot obtain estimates for the raw maximum HR-QoL loss in the confirmed cases. Therefore, using the responses from the suspected cases, we used a mixture model approach to estimate the maximum HR-QoL loss as a function the independent variables—age (5–14 years, 15 years and older), coughing severity, healthcare seeking behaviour, productivity and VAS score loss. We classified the adjusted maximum HR-QoL loss for each RSV episode as either severe (above a threshold value h) or mild (equal to or below h), with probability p and 1-p respectively. For suspected cases with a raw maximum HR-QoL loss above *h*, we used a linear regression model to estimate the adjusted maximum HR-QoL loss in severe episodes as a function of the independent variables. Similarly, for suspected cases with a raw maximum HR-QoL loss equal to or below *h*, we used a log-transformed linear regression model to estimate the adjusted maximum HR-QoL loss in mild episodes as a function of the independent variables. The raw maximum HR-QoL loss values were log-transformed to prevent negative adjusted maximum HR-QoL loss values. Finally, by transforming the raw maximum HR-QoL loss to a binary variable—0 for mild infections and 1 for severe infection—we used a logistic regression to calculate the probability of a severe episode p as a function of the independent variables. The final three regression models contain those variables that significantly influenced the adjusted maximum HR-QoL at the 5% significant level (Appendix II). The variance of the full mixture model was estimated by simulation using prediction intervals for the linear and log-transform linear model and confidence intervals for the logistic regression model. The threshold value h was estimated by finding the minima of the fitted mixture distribution for the raw maximum HR-QoL loss values. To assess the accuracy of the model, we compared the distributions derived from between the raw and adjusted maximum HR-QoL loss for the suspected cases for each of the independent variables considered in the model calibration. The regression analysis, hypothesis testing for significant covariates, and stochastic simulation was performed in R (v. 3.3.2), and plotting was performed in Mathematica (v. 10.3.0.0).

### Quality-adjusted life year (QALY) loss due to an RSV episode

For confirmed cases in children under the age of five years and suspected cases in persons aged five years and older, we estimated each respondent’s QALY loss by multiplying the adjusted maximum HR-QoL life loss by the duration of coughing and a scaling factor for disease severity throughout this duration. We estimated the distribution of coughing duration for confirmed and suspected cases separately by pooling responses, respectively (Appendix II). We estimated the scaling factor for disease severity using detailed data from the Flu Watch study,^13^ Flu Watch is a community cohort study in which householders were asked to record all respiratory infections and submit a nasal swab for Polymerase Chain Reaction (PCR) based identification of respiratory viruses over winter seasons. In 2010/11 participants or adult carers were also asked to complete daily EQ-5D instruments during each day of illness. These five sets of responses resulted in daily HR-QoL estimates from five community cases of confirmed RSV (ages 16–45), from which we quantified five estimates of the scaling factor of disease severity as the average daily severity of the symptoms relative to the maximum HR-QoL loss. We then took the mean of these values to estimate the scaling factor for disease severity. It is not possible to obtain estimates for the QALY loss in confirmed infections who are under the age of five years who did not seek healthcare as the confirmed cases were recruited into the study conditional on them seeking healthcare. Therefore, to gain QALY loss estimates we assume that the ratio of QALY loss for people over five years who seek healthcare to those that do not is the same independent of age.

### Total QALY loss due to RSV for confirmed infections

We estimated the annual QALY loss due to RSV infections in the UK for healthcare seeking RSV episodes in children under five years old and for individuals five years and older by multiplying the respective QALY loss per episode calculated in this study with a previous estimate of the respective age-specific annual number of GP consultations and hospital admissions due to RSV in the UK.^14,15^

### Ethics approval

In accordance with The Health Service (Control of Patient Information) Regulations 2002 No. 1438 Section 251 Regulation 3. Public Health England may process confidential patient information with a view to monitoring and managing; outbreaks of communicable disease; incidents of exposure to communicable disease and the delivery, efficacy, and safety of immunisation programmes.^16^ All questionnaires were returned from households and stored at PHE were anonymised.

## Results

### Questionnaire responses

We sent out 770 questionnaire packs between 15 December 2016 and 4 January 2017 and received 122 responses by 28 February 2017 (response rate of 16%). We found that, when stratified by year of age, the age distribution of the confirmed cases who responded was similar to the age distribution of the contacted confirmed RSV index cases. However, when stratified by month of age in the first year of life, we oversampled infants 3–4 months old and undersampled infants 1–2 month old (**Figure 1a and b**). In the 122 households, suspected cases were reported in 33 (27·0%) persons aged 5–14 years old and 54 (44·2%) of persons aged 15 years or older.

**Figure 1:**
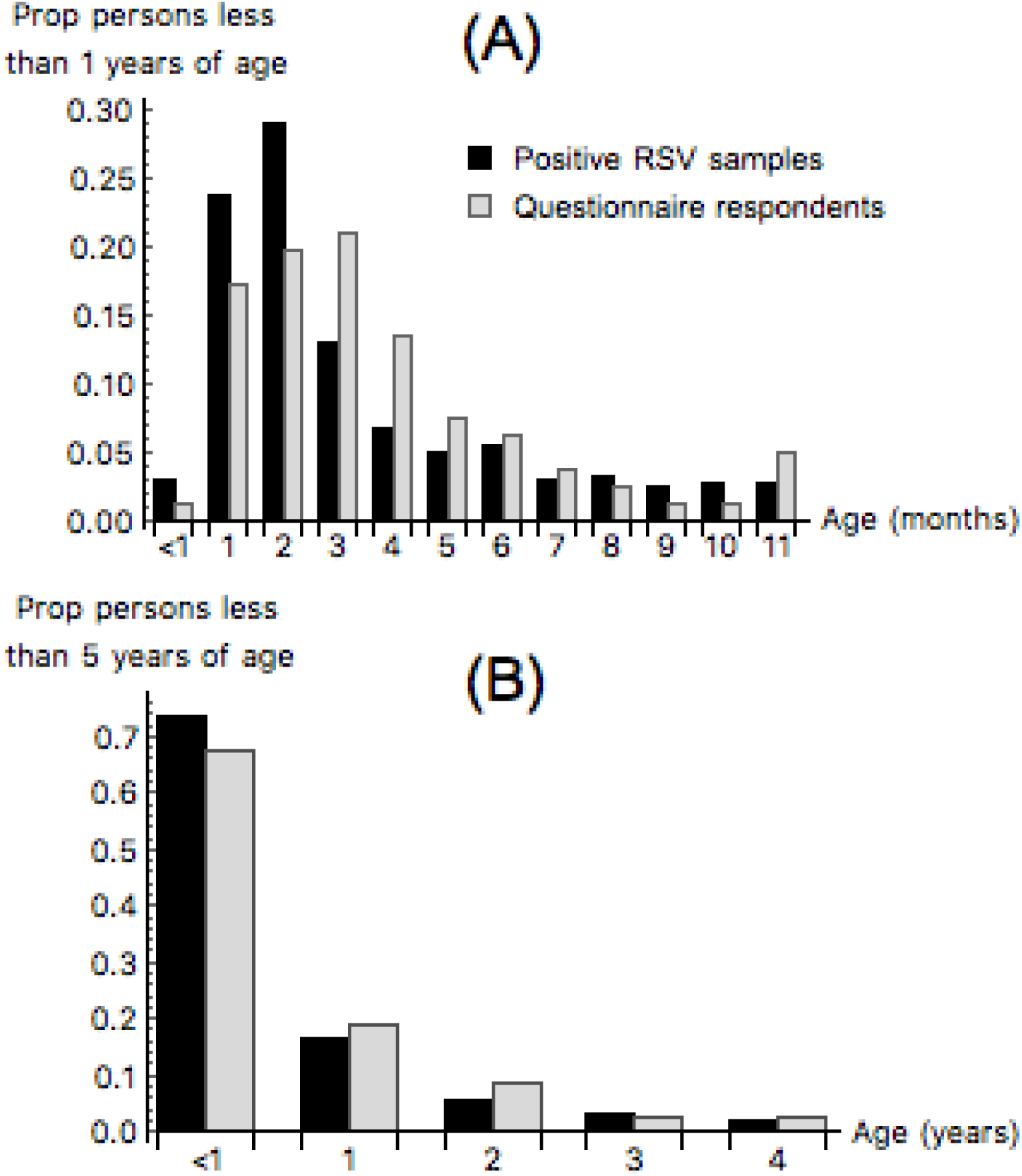
Age of confirmed RSV samples in PHE database (N=770, black) and of returned for analysis (N=122, gray).

In suspected cases, 25/33 (75·8%) of 5–14-year-olds and 48/54 (88·9%) of respondents aged 15 years and older completed all questions in the EQ-5D-3L and EQ-5D-Y instruments, respectively, to allow the calculation of the raw maximum HR-QoL weight. To estimate the adjusted maximum HR-QoL loss, we included all questionnaires from 5-14 years old and person 15 years and older (21/33 (63·6%) and 40/58 (69·0%) respectively) which, in addition to completing all five dimensions of health status in EQ-5D instruments, completed answers for age, productivity loss, healthcare-seeking behaviour, coughing severity and VAS score loss. To estimate the adjusted maximum HR-QoL loss in the confirmed cases, we used 108/122 (88·5%) of the questionnaires which provided the answers to coughing severity, productivity loss, healthcare-seeking behaviour, and VAS score loss. Duration of coughing was provided for 98/122 (80·3%) and 43/87 (49·4%) of confirmed and suspected cases respectively.

In the questionnaire responses from the suspected cases, we found that 21/33 (63·6%) of children aged 5-14 years old, and 43/54 (79·9%) of persons aged 15 years and older did not seek any healthcare because of their suspected RSV episode (**Table 1**). Further, we found that 17/33 (51·5%) children aged 5–14 years old took time off school and 9/54 (16·6%) of persons aged 15 years and older took time off work or school due to their suspected RSV infection, both with a median time off of 2 days (range 1–10) (**Table 1**). The EQ-5D-Y instruments suggested that for children aged 5-14 years old the dimensions of healthcare which were most affected by RSV were the effect on usual activities (72%), pain/discomfort (76%) and anxiety/depression (84%). The EQ-5D-3L responses for respondents aged 15 years and older suggested similar results with RSV affecting respondents’ usual activities (54·2%), causing pain/discomfort (36·0%) and anxiety and depression (32·0%) (**Figure 2**). After using the UK TTO scoring tariff, the raw HR-QoL weight for the worst day of infection for children aged 5–14 years old and persons 15 years and older was 0·630 (range -0·429–1·000) and 0·630 (range -0·429–1·000) respectively. (**Table S7**). This led to a raw maximum HR-QoL loss of 0·456 (range 0·0–1·170) and 0·358 (range 0–0·998) for 5-14 and 15 years and older respectively. As there was no significant difference between the raw maximum HR-QoL loss between 5–14 years old and persons 15 years (Kolmogorov-Smirnov test, p = 0·291), all HR-QoL and QALY results were pooled for further analysis.

**Table 1.** Summary of index, 5-14, and 15+ Questionnaire responses· Numbers in parentheses is the percentage unless otherwise stated. VAS, visual analogue scale.

|  | * Aged 0–4 years (n = 122) (%) | Aged 5–14 years (n = 33) (%) | Aged 15 years and over (n = 54) (%) |
| --- | --- | --- | --- |
| **Symptoms** |  |  |  |
| Runny/blocked nose | 96 (78.7) | 28 (84.8) | 43 (79.6) |
| Fever | 70 (57.4) | 18 (54.5) | 22 (40.7) |
| Coughing | 110 (90.2) | 27 (81.8) | 51 (94.4) |
| Sore throat | 36 (29.5) | 17 (51.5) | 38 (70.4) |
| **Coughing severity** |  |  |  |
| No effect on daily activities | 16 (13.1) | 10 (30.3) | 8 (14.8) |
| Mild effect on daily activities | 34 (27.9) | 15 (45.5) | 32 (59.3) |
| Severe effect on daily activities | 93 (76.2) | 4 (12.1) | 8 (14.8) |
| **Coughing severity duration** |  |  |  |
| No effect on daily activities (median, range) | 4 days (1–14) | 10 days (10–10) | 14.5 days (1–28) |
| Mild effect on daily activities (median, range) | 3.5 days (1–14) | 3 days (1–9) | 5.5 days (1–28) |
| Severe effect on daily activities (median, range) | 6.5 days (1–35) | 3 days (1–4) | 7 days (3–10) |
| **Healthcare seeking** |  |  |  |
| Phone/email NHS 111 / NHS 24 / NHS Choices | 39 (32.0) | 2 (6.1) | 2 (3.7) |
| Phone/email GP—response from the receptionist | 20 (16.4) | 2 (6.1) | 2 (3.7) |
| Phone/email GP—response from the doctor or nurse | 20 (16·4) | 2 (6·1) | 2 (3·7) |
| Visit a GP or nurse | 83 (68·0) | 10 (30·3) | 10 (18·5) |
| Visit A&E department (including out of hours service) | 71 (58·2) | 3 (9·1) | 1 (1·9) |
| Admitted to hospital | 103 (84·4) | 1 (3·0) | 1 (1·9) |
| None | 0 (0·0) | 21 (63·6) | 43 (79·6) |
| **Productivity** |  |  |  |
| Individuals reporting taking time off work or school | — | 17 (51·5) | 9 (16·7) |
| Duration of time off work or school (median, range) | — | 2 days (1–10) | 2 days (1–7) |
| **VAS score** |  |  |  |
| Baseline (median, range) | 90 (30–100) | 95 (10–100) | 95 (50–100) |
| Worst day (median, range) | 20 (0–85) | 50 (5–85) | 50 (0–90) |
| Loss (median, range) | 65 (10–100) | 38 (0–90) | 35 (10–85) |

\* Conditional on ascertaining a confirmed case through GP/hospitalisation.

**Figure 2:**
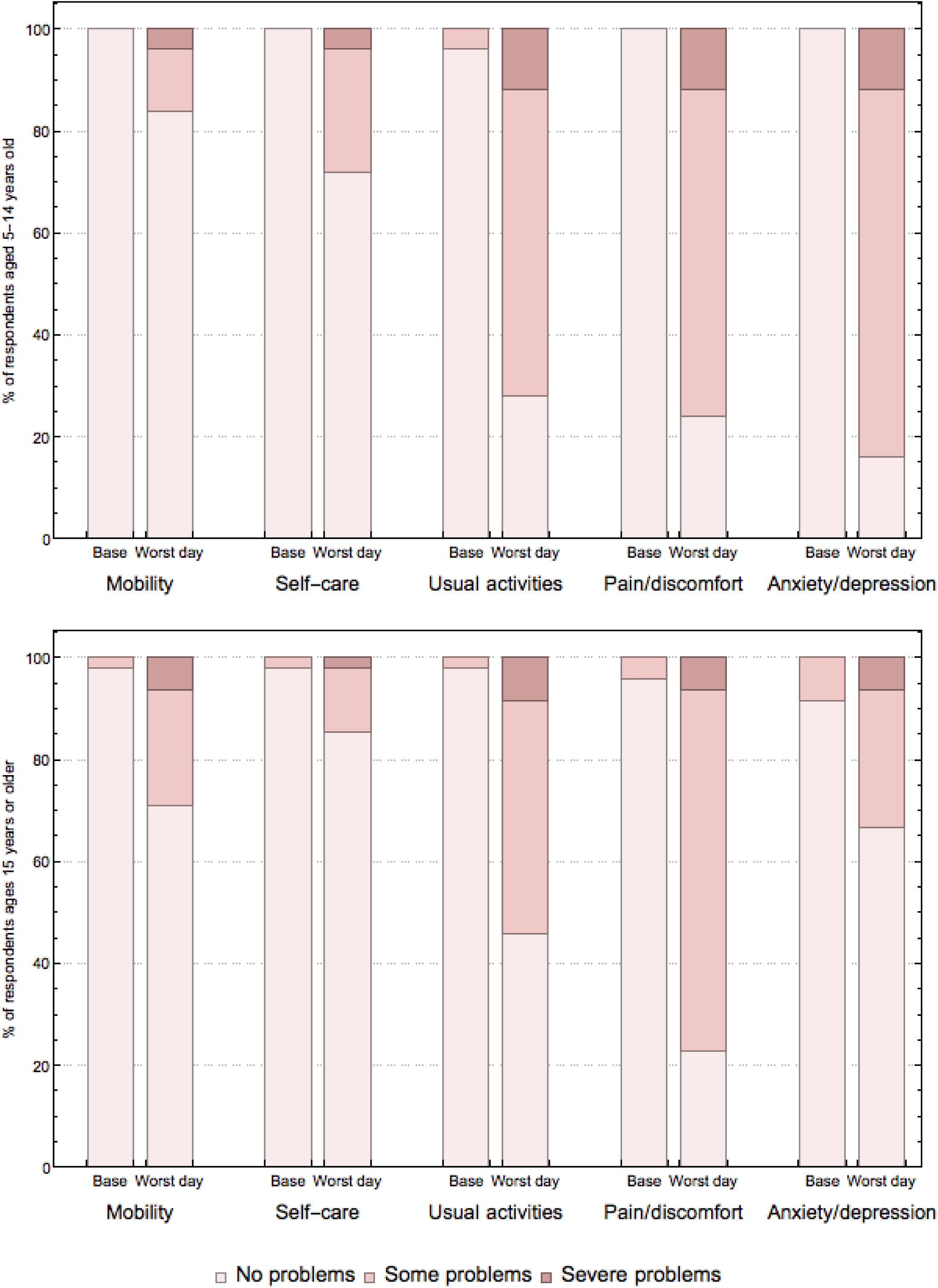
Responses from the EQ-5D-3L-Y and EQ-5D-3L instruments on the base and worst day of health for respondents aged 5-14 years old (top) and 15+ years old (bottom).

### Adjusted maximum HR-QoL loss

We found the threshold value for the maximum HR-QoL loss, above which infections are considered severe, to be h = 0·582 (**Figure S1**). Using linear regression, we found that VAS score loss significantly adjusted the adjusted maximum HR-QoL loss for mild infections (p < 0·001), and healthcare-seeking behaviour (none or any) for severe infection (p = 0·0012). Using logistic regression, we found that coughing severity (none/mild or severe) significantly adjusted the probability of severe infections (p < 0·001) (**Table S2 and Figures S3–S6**).

Our full mixture model estimated the adjusted maximum HR-QoL loss in suspected cases aged five years and older who did and did not seek health care as 0·616 (95% CI 0·155–1·371) and 0·405 (95% CI 0·111–1·137) respectively. We used the calibrated mixture model to estimate the adjusted maximum HR-QoL loss for confirmed cases less than five years of age, who sought health care as 0·820 (95% CI 0·222–1·450). In assessing the model accuracy, we found the difference between the means of the HR-QoL loss between the raw and adjusted maximum HR-QoL loss for each of the independent variables was 0·063 (range 0·026–0·154) (**Figure 3**).

**Figure 3:**
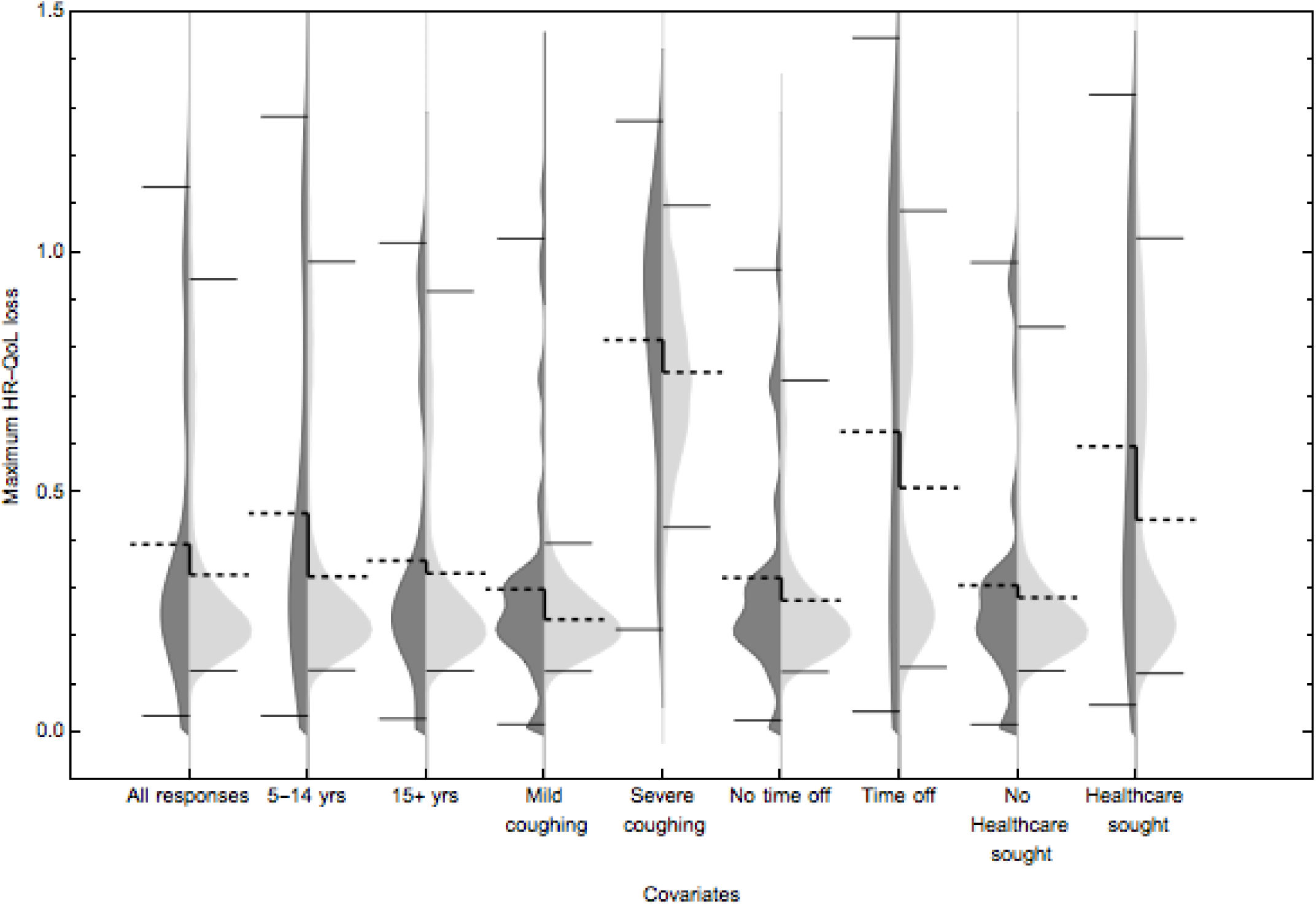
Distributions of the maximum HR-QoL loss from the EQ-5D instruments (dark gray) and estimated using statistical model (light gray.) The dashed line shows the mean, and solid thin lines indicate the upper and lower 95% CI.

### Quality-Adjusted Life Year loss

The duration of coughing in children under five years old (median 7 days (range 1–22)) was longer than in those aged five years and older (median 4 days (range 1–20)) (**Figure S7**). The daily RSV HR-QoL weights from Flu Watch^13^ suggest that for half of the duration of symptoms, the HR-QoL weight decreases linearly to the minimum HR-QOL weight, and then linearly increases to baseline health. We thus estimated the mean scaling factor for disease severity as 0·25. By calculating QALY loss as the product of the adjusted maximum HR-QoL loss, duration of coughing and the scaling factor for disease severity we quantified the QALY loss for suspected episodes in persons aged five years and older as 1·950 × 10^−3^ (95% CI (0·185–9·578 × 10^−3^) for those who seek healthcare and 1·543 × 10^−3^ (95% CI 0·136–6·406 × 10^−3^) for those who do not seek healthcare. For confirmed cases in children less than five years old, all of whom sought healthcare, the QALY loss per episode was 3·823 × 10^−3^ (0·492–12·766 × 10^−3^), (**Tables 2, S8, S9**).

**Table 2.** HR-QoL, QALD, and QALY loss for significant factors in the confirmed cases in children less than five years of age, and in the suspected cases in children five years and older.

|  | Under five years of age* | Five years of age and older |
| --- | --- | --- |
| **Mean maximum HR-QoL loss (95% CI)** |  |  |
| *Coughing severity* |  |  |
| None or mild | 0.499 (0.148–1.482) | 0.382 (0.111–1.113) |
| Severe | 0.878 (0.344–1.443) | 0.785 (0.280–1.368) |
| *Healthcare seeking behaviour* |  |  |
| None | 0.539 (0.144–0.952)** | 0.405 (0.111–1.137) |
| Seek healthcare | 0.820 (0.222–1.450) | 0.616 (0.155–1.371) |
| **Mean QALD loss (95% CI)** |  |  |
| *Coughing severity* |  |  |
| None or mild | 0.845 (0.097–3.292) | 0.528 (0.050–2.167) |
| Severe | 1.496 (0.221–4.841) | 1.103 (0.126–4.149) |
| *Healthcare seeking behaviour* |  |  |
| None | 1.00 (0.141–3.652)** | 0.565 (0.049–2.349) |
| Seek healthcare | 1.391 (0.179–4.617) | 0.866 (0.071–3.508) |
| **Mean QALY loss (95% CI)** |  |  |
| *Coughing severity* |  |  |
| None or mild | $2.336 \times 10^{-3}$ (0.269–9.255) | $1.448 \times 10^{-3}$ (0.135–5.928) |
| Severe | $4.098 \times 10^{-3}$ (0.624–13.141) | $2.990 \times 10^{-3}$ (0.346–11.387) |
| *Healthcare seeking behaviour* |  |  |
| None | $3.024 \times 10^{-3}$ (0.329–10.098)** | $1.543 \times 10^{-3}$ (0.136–6.406) |
| Seek healthcare | $3.823 \times 10^{-3}$ (0.492–12.766) | $1.950 \times 10^{-3}$ (0.185–9.578) |

\*Conditional on ascertaining a confirmed case through GP/hospitalisation.
\*\*Implicitly estimated by assuming the proportional reduction in HR-QoL loss and QALY loss is the same as observed in suspected infections in persons over five years of age.

### Implications for economic evaluations

Previous studies have estimated the combined number of annual GP consultations and hospital admissions due to RSV in the UK as 110,016 (range 62,414–157,617) for children aged 5–17 years and 505,046 (range 336,305–604,873) for adults.^14,15^ Using our QALY loss estimate of 1·950 × 10^−3^ for healthcare seeking episodes, this results in a mean annual loss of 1,199 (range 777–1487) QALYs for healthcare seeking infections in individuals five years and older in the UK. Similarly, for children under five years, the estimated number of both GP consultations and hospital admissions due to RSV in the UK is 369,302 (range 253,825–467,277).^15^ Using our estimate of 3·823 × 10^−3^ for the QALY loss for healthcare seeking episodes in children under the age of five years, this results in a mean annual QALY loss of 1,411 (range 970–1786). These results suggest that 45·9% of the annual QALY loss due to RSV episodes seeking healthcare is attributable to persons five years and older.

Further, using our result that 25% of individuals aged five years and older seek healthcare, we estimate that there are approximately 1.8 (range 1.6–3.0) million RSV infections in the UK annually that will not be captured in a healthcare-focussed surveillance system. The mean annual QALY loss associated with these non-surveilled episodes for persons five years and older is around 2,900 (range 2,460–4,706), suggesting that approximately 29% of the QALY loss in this age group can only be captured by community surveillance.

## Discussion

In this study, we quantified the quality of life (QALY) loss associated with RSV episodes. For children over five years old and adults, we found that the QALY loss can be accurately predicted by whether there was severe coughing, whether healthcare was sought, and Visual Analogue Scale score loss. We used a novel statistical model to evaluate the QALY loss in children under five years old, in whom the majority of severe RSV episodes occur. We found the QALY loss in children under the age of five years who sought healthcare is 3·823 × 10^−3^ (95% CI 0·492– 12·766), and for persons five years and older to be 1·950 × 10^−3^ (95% CI 0·185–9.578) for those who seek healthcare and 1·543 × 10^−3^ (95% CI 0·136–6·406) and for those who do not seek healthcare. In addition, we found the 73·6% of infections in persons over the age of five did not seek healthcare, and 30·0% took time off due to their infection.

Our study has some limitations. First, because the confirmed cases were recruited into the study conditional on them seeking healthcare, we could not estimate the QALY loss in children less than five years old who did not seek healthcare. To overcome this limitation, we assume that the ratio of QALY loss for people over five years who seek healthcare to those that do not is the same independent of age. To collect data directly on the QALY loss in children under five who do not seek healthcare would require a much larger and more intensive community-based study with frequent testing throughout an RSV season. Second, suspected cases may have experienced non-RSV respiratory disease. However, as previous studies have shown that around 70% of households experience a second infection in either siblings or parents during the same time as an infant, we think it is reasonable to assume that the majority of suspected cases are in fact RSV.^17,18^ Finally, completing questionnaires some days after symptoms may be subject to recall bias as our estimates for the maximum HR-QoL life loss for persons aged 15 years and older (0.452 (95% CI 0.177–1.222)) are larger to the maximum HR-QoL loss estimated during the infection in the Flu Watch study (range 0.107–0.309).

Our study is the first to estimate the QALY loss due to acute RSV infection across all ages. In addition, we developed a novel method to estimate QALY loss due to RSV in young children for whom standardised instruments for deriving HR-QoL estimates are not appropriate. Thus our method leverages the use of standardised instruments such as EQ-5D to quantify QALY loss using more easily measurable variables of infection in young children. In particular, we derived a method for determining the maximum HR-QoL loss due to RSV infection. We found two previous studies which also estimated the HR-QoL due to RSV infection both of which suffer from shortcomings. The first is a Time Trade-off study which derived HR-QoL life estimates using responses from participants about a hypothetical illness that they, or their child, had not experienced. ^19,20^ Our study is advantageous because the adjusted maximum HR-QOL loss values are the subjective HR-QoL estimates from people who have, or suspected to have experienced an RSV infection. The second study estimates the HR-QoL using EQ-5D instruments, however the results are derived by describing the current condition of a five-year-old child with a history of RSV—it is not a health utility for an RSV episode. Therefore, this estimate is for RSV sequelae, which can complement our estimate for RSV infections derived in this study. Despite the limitations in this study, it is the primary source of HR-QoL estimates for existing RSV cost-effectiveness analysis.^19^ Using HR-QoL loss for RSV sequelae of acute RSV infection in cost-effectiveness analysis could underestimate the impact of potential intervention strategies. We recommend that future cost-effectiveness analyses additionally use directly obtained HR-QoL loss estimates for RSV episodes, such as those presented in this paper, to ensure that the cost-effectiveness of potential intervention programmes is accurately determined.

We compared the adjusted maximum HR-QoL loss found in the novel statistical model with the raw maximum HR-QoL loss for independent variables used to calibrate the transmission model to assess the model accuracy. We found that the difference between the means of the two methods of calculating the maximum HR-QoL was less than 0·1 HR-QoL loss for 66% of the independent variables considered, implying that the model predicts with accuracy.

We estimated a QALY loss of 1·758 × 10^−3^ (95% CI 0·150–7·303 × 10^−3^) due to a suspected RSV episode in persons aged five years and older. Comparing our study to the Flu Watch prospective cohort study, we found that the QALY loss for an RSV episode is less than the QALY loss for a confirmed Influenza H1N1 episode, which is estimated to have mean 4·4 × 10^−3^ (range -2·5–18·2 × 10^−3^) across all ages.^22^ However, our estimates are similar to the QALY loss for cases of respiratory disease in the same study which were not confirmed to be Influenza, but were confirmed to suffer from a fever and reported coughing/sore throat (mean 2·6 × 10^−3^ (range -69·2–39·7 × 10^−3^) across all ages). We cannot compare the QALY loss due to the confirmed RSV episodes in children under the age of five (3·823 × 10^−3^ (95% CI 0·492–12·766 × 10^−3^)), with values from the Flu Watch because the confirmed RSV infections were recruited dependent on an infection being severe and requiring medical attention. We therefore compare estimates to a different study which suggests that these RSV severe episodes are milder than hospitalised Influenza episodes, (QALY loss of 6·0 × 10^−3^ (range 5·1–6·9 × 10^−3^)) but similar in severity to non-Influenza episodes who suffer Influenza-like illness (ILI) and present at a clinical interface (4·0 × 10^−3^ (range 3·4–4·6 × 10^−3^)).^23^ These comparisons suggest that, though Influenza has a higher QALY loss per episode, the QALY loss due to an RSV episode is comparable to previous QALY loss estimates for persons with general ILI. In a wider context, comparing with QALY estimates derived via EQ-5D instruments with UK TTO scoring tariff for measles, another disease with high burden in children, the QALY loss was 0·019 (95% CI 0·016–0·022) per episode.^24^

We estimated that 46% of the QALY loss associated with healthcare seeking episodes was attributable to individuals five years and older. This result suggests that neglecting QALY loss in older children and working age adults might substantially underestimate the impact of a potential RSV vaccine programme. Further, RSV is characterised by high levels of household transmission,^25,26^ which points to a need to evaluate both the direct and herd protection effects of potential vaccination strategies into impact assessments. Together, this evidence suggests that integrating transmission models into economic evaluations will be important to accurately estimate the impact of potential vaccine programmes across the entire population.

In this study we are only able to directly estimate the QALY loss for children under five years old who do not seek healthcare, we cannot evaluate the frequency at which this occurs. However, we expect the probability of healthcare seeking in children under the age of five years to be higher than the 25% reported in those aged five years and older for two reasons: because infections in infants are generally more severe, with higher rates of symptomatic infections,^27^ and because of possible increased parental healthcare seeking for infants.^28^ Regardless, the healthcare seeking behaviour for both children and adults will likely depend on the region and careful consideration will need to be taken into account in economic evaluations.

In summary, we estimated the QALY loss due to an RSV episode in confirmed cases in children less than five years old and suspected cases in persons five years old or older. The QALY loss due to an RSV episode is less than the QALY loss due to an influenza episode. In addition, RSV infections in individuals aged five years and older account for 46% of the annual QALY loss attributable to healthcare seeking episodes in the UK. Consequently, economic evaluations of potential vaccine programmes should consider the effect on reducing incidence not only where the severe disease burden lies, but across the across the whole population.

## Acknowledgements

KEA acknowledges funding from the National Institute for Health Research Health Protection Research Unit in Immunisation at the London School of Hygiene and Tropical Medicine in partnership with Public Health England. DH received funding from a Medical Research Council PhD Studentship (administered through CoMPLEX University College London). DT received funding from the European Union’s Horizon 2020 research and innovation programme under grant agreement #634446 for the I-MOVE+ (Integrated MoGnitoring of Vaccines in Europe) project. JPG’s research is supported by the National Institute for Health Research (NIHR) Collaboration for Leadership in Applied Health Research and Care North Thames at Bart’s Health NHS Trust (NIHR CLAHRC North Thames). The views expressed are those of the authors and not necessarily those of the National Health Service, the Medical Research Council, National Institute for Health Research, Department of Health, European Union, or Public Health England. We thank Nick Andrews for helpful advice and guidance with the statistical analyses.

## Author’s contributions

DH, KEA, MB, JPG, AT, AJvH, RP conceived and designed the surveillance study. DH, DT, and HZ were involved in the data extraction, the distribution of the questionnaires, and data input of the questionnaire responses. EF and AH were involved in collecting and interpreting the Flu Watch data. DH performed the statistical analysis with interpretations from KEA, MB, JPG, RP. DH, KA, RP, MB, JPG drafted the manuscript with critical revisions from DT, AJvH, HZ, EF, AH.

